# Fructose Metabolism Contributes to the Warburg effect

**DOI:** 10.1101/2020.06.04.132902

**Authors:** Bing Han, Lu Wang, Jingyu Zhang, Meilin Wei, Cynthia Rajani, Runming Wei, Jingye Wang, Haining Yang, Michele Carbone, Guoxiang Xie, Wen Zhou, Wei Jia

## Abstract

Fructose metabolism is increasingly recognized as a preferred energy source for cancer cell proliferation. However, it remains unclear why cancer cells favor fructose metabolism and how they acquire a sufficient amount of fructose. Here we report that cancer cells convert glucose into fructose through intra- and extracellular polyol pathways. The fructose metabolism bypasses normal aerobic respiration’s self-control to supply excessive metabolites to glycolysis and promotes the Warburg effect. Inhibition of fructose production drastically suppressed glycolysis and ATP production in cancer. Furthermore, we determined that a glucose transporter, SLC2A8/GLUT8, exports intracellular fructose to other cells in the tumor microenvironment. Taken together, our study suggests that the Warburg effect is achieved by means of fructose metabolism, instead of glucose metabolism alone.

## INTRODUCTION

Cancer cells consume excessive amounts of glucose to sustain their unrestricted proliferation(Goncalves et al., 2019). We previously reported that cancer cells such as acute myeloid leukemia cells exhibited enhanced fructose utilization under low glucose conditions and that pharmacologic blockade of fructose utilization suppressed AML cell proliferation and enhanced the therapeutic efficacy of Ara-C(Chen et al., 2016). Our recent findings, along with reports from other groups(Chen et al., 2020; Chen et al., 2016; Goncalves et al., 2019; Krause and Wegner, 2020), suggest that fructose is a preferred carbon source for many cancer cells. High expression of the fructose transporter SLC2A5/GLUT5 was found in multiple types of cancers, enabling increased uptake of fructose from the microenvironment and enhanced cancer growth and development. However, dietary fructose is mostly converted to glucose, glycogen, and organic acids in the intestine and liver(Jang et al., 2018). Therefore, the fasting blood levels of fructose are about 1,000-fold lower (∼0.005 mM) than glucose (∼5.5 mM) under normal physiological conditions. We thus hypothesized that this unique metabolic feature in cancer cells must arise from additional fructose sources and that increased fructose utilization conferred advantages over the use of glucose.

The polyol pathway is commonly present in somatic cells. It converts about 3% of intracellular glucose to fructose under normal physiological conditions in a two-step process(Yabe-Nishimura, 1998). The first step involves reducing glucose to sorbitol utilizing the reducing equivalents from NADPH and the aldose reductase enzyme, AKR1B1. The second step is the oxidation of sorbitol to fructose via sorbitol dehydrogenase (SORD) and the conversion of NAD^+^ to NADH.

Dr. Otto Heinrich Warburg first observed that cancer cells tend to ferment glucose to lactic acid even in aerobic conditions, a phenomenon later known as “aerobic glycolysis, or the Warburg effect”. The current understanding of this phenomenon is that glucose fermentation promotes biomass synthesis in cancers to maintain cell proliferation(Vander Heiden et al., 2009). The Warburg effect has been extensively studied and also found in non-cancerous, high-proliferating cells, such as T lymphocytes(Abdel-Haleem et al., 2017), however its precise metabolic mechanism remains unclear.

## RESULTS

### Intra and extracellular polyol pathways supply fructose to cancer cell metabolism

This study revealed non-dietary sources of fructose produced by intra- and extracellular polyol pathways due to cancer-mediated metabolic reprogramming. First, we identified significant levels of ^13^C-fructose in 11 out of 12 cancer cell lines that were incubated with ^13^C-glucose (Figure 1A). Basal levels of fructose were also detected in the two control cell lines, the pancreatic epithelial cell, HPDE6C7, and the mammary epithelial cell line, MCF-10A. We then tested the fructose-conversion rate in the A549 cell line. The results showed that A549 cells converted glucose to fructose rapidly and that their intracellular ^13^C-fructose level reached a saturation point after 2 hours of treatment with 25 mM ^13^C-glucose. We also found that the intracellular ^13^C-fructose level was equal to intracellular ^13^C-glucose after 2 hours (Figure 1B). Therefore, we estimated that about 50% of the glucose taken up by A549 cells was converted to fructose.

**Figure 1.**
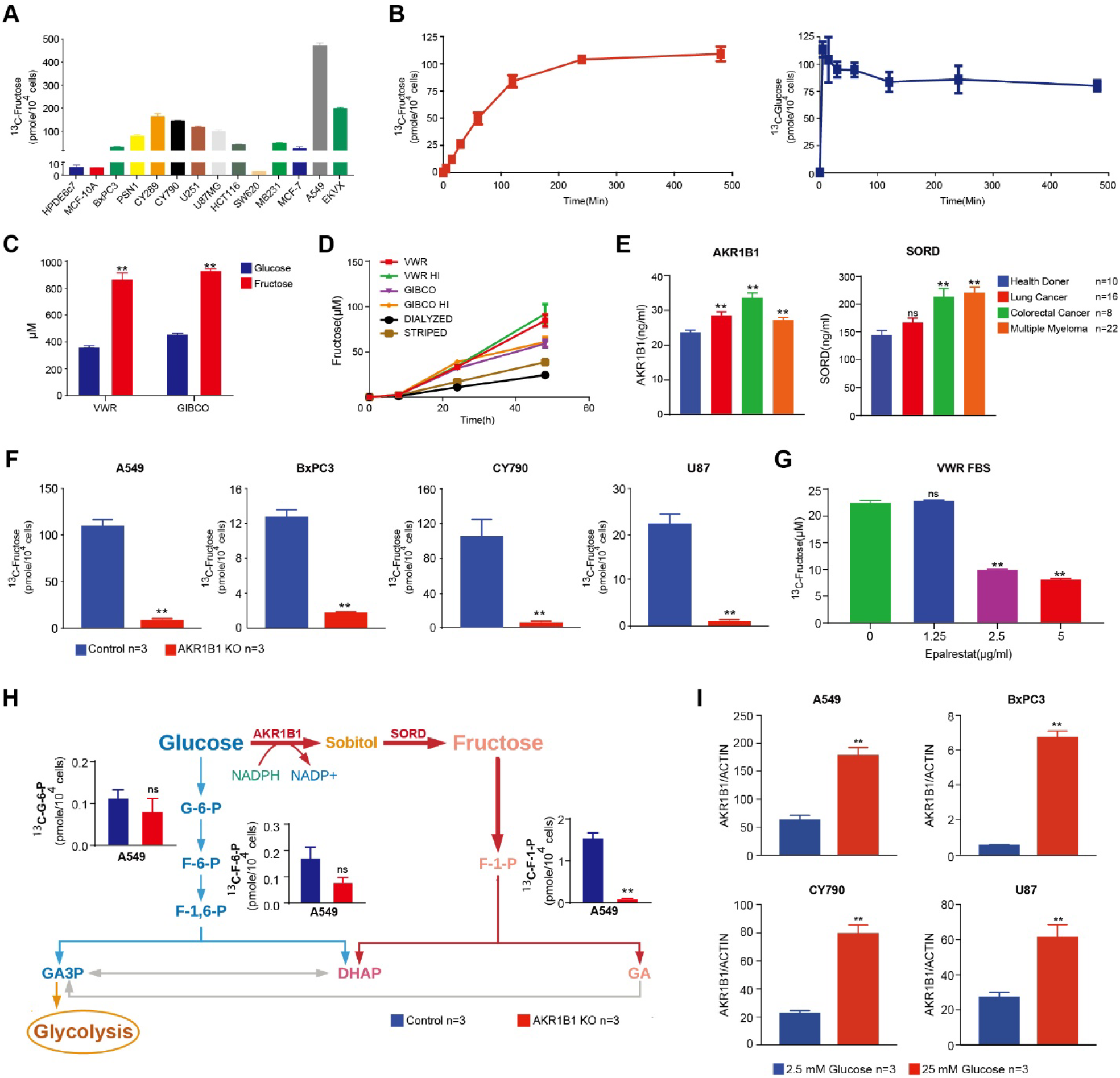
Cancer cells produce high concentrations of fructose. **(A**) HPDE6C7 and MCF-10A cells are considered normal cells controls. Along with these controls, we tested a wide array of different cancer cell lines using ^13^C labeled metabolic flux analysis. From these data, it was seen that although most of the cells affected the conversion of glucose to fructose, there was variability in how much fructose was produced. Notably, A549 cells produced the most fructose. **(B)** Based on the results in Figure 1a, we tested the rate of fructose production in A549 cells. The red line represents intracellular ^13^C-fructose production, and the blue line represents the intracellular ^13^C-glucose level. **(C**) The concentration of glucose and fructose in two major commercial FBSs. (**D**) The ^13^C-fructose production rate in 6 different FBSs. (**E**) Amounts of AKR1B1 and SORD in sera of human cancer patients. (**F**) AKR1B1 KO blocked fructose production in tested cancer cells. (**G**) AKR1B1 inhibitor, epalrestat, suppressed the ^13^C-fructose transformation in FBS. **(H)** Sketch of the polyol pathway and fructose metabolism. In A549 cells, fructose specific metabolite, fructose-1-phosphate (F-1-P), highly depended on the polyol pathway. Only a trace amount of F-1-P can be detected in AKR1B1 KO cancer cells, while the compromised polyol pathway did not significantly affect glucose metabolism. The amounts of glucose specific metabolites, glucose-6-phosphate and fructose-6-phosphate (G-6-P and F-6-P), had no significant changes in A549 AKR1B1 KO cancer cells. (**I**) We tested the regulation of expressions of AKR1B1 according to glucose level in different cancer cells. The AKR1B1 expression was promoted significantly by high glucose level (25mM) in all tested cell lines. Data are represented as mean ± SEM. *: t-test p<0.05; **: t-test p<0.01.

In addition to intracellular production of fructose, we unexpectedly found that in two common commercial brands of FBS, there were high concentrations (approximately a double amount) of fructose relative to glucose (Figure 1C). To identify the possible reasons for high fructose levels in FBS, we selected regular, heat-inactivated (HI), active charcoal striped, and dialyzed FBSs purchased from two major commercial vendors (VWR and Gibco) for this study. We adjusted every FBS sample to 25mM of ^13^C-glucose and cultured them at 37°C, 5% CO_2_ for 0, 8, 24, and 48 hours. The ^13^C-fructose levels increased with culture time in all the FBS samples and different conversion rates were observed depending on the type of FBS sample (Figure 1D). These results revealed a novel source of fructose: a mechanism that naturally converts glucose to fructose in blood.

The polyol pathway is the only known physiological reaction to convert glucose to fructose intracellularly. We analyzed two enzymes involved in the polyol pathway, AKR1B1 and SORD, at both gene (Figure S1A) and protein (Figure S1B) levels and found that all cancer and control cell lines expressed varying levels of these two genes. We then examined the human TCGA pan-cancer database (Figure S1C) and confirmed that all human cancers express AKR1B1 and SORD, which suggested that the polyol pathway was active as an essential metabolic feature in all types of cancer. To identify whether the polyol pathway also processes extracellular glucose-fructose conversion, we then assayed the AKR1B1 and SORD proteins in FBS by Western Blot (Figure S1D). Both of these two components were found in all tested samples with varying amounts. Additionally, we measured the concentrations of AKR1B1 and SORD in sera of 10 healthy donors, 16 lung cancer, 8 colorectal cancer, and 22 multiple myeloma patients. Serum levels of AKR1B1 were significantly higher in all cancer patients than healthy controls, while serum levels of SORD were significantly increased in colorectal cancer and multiple myeloma patients, but not in lung cancer (Figure 1E). We then blocked the intracellular and extracellular polyol pathway by AKR1B1 gene knocked out (KO) (Figure S1E) or epalrestat (Goto et al., 1995), an AKR1B1 inhibitor, respectively, to verify the role of the polyol pathway in glucose-fructose conversion. Inhibition of AKR1B1 by either method effectively decreased the intracellular (Figure 1F) and extracellular (Figure 1G) fructose concentrations. These results provide evidence for the activity of the polyol pathway in cancer cells and serum.

As a result of AKR1B1 gene KO in cancer cells, the fructose specific metabolite, fructose-1-phosphate (F-1-P), was significantly decreased as shown in Figure 1h and Figure S1f. Inhibition of the polyol pathway did not have the same effect on the production of the two glucose-specific metabolites, glucose-6-phosphate (G-6-P) and fructose-6-phosphate (F-6-P), in the four cancer cell lines (Figure 1H and Figure S1G). Therefore, AKR1B1 KO blocked ^13^C-fructose production from the polyol pathway but did not alter glucose metabolism. Thus, it appeared that conversion from fructose to glucose is insignificant, and that once fructose is produced in cancer cells, its subsequent metabolism is independent of glucose metabolism. Fructose-specific metabolism is different from glucose metabolism in that it has fewer steps, and the pathway is not reversible (Sun and Empie, 2012). Cancer cells commonly overexpress glucose transporters, such as SLC2A1/GLUT1 and SLC2A3/GLUT3, to take in a large amount of extracellular glucose to support their high rates of proliferation (Krzeslak et al., 2012). Glucose is known to be the preferred carbon source for the production of structural components necessary for cell growth and division. When cancer cells metabolize glucose to synthesize biomass, a massive amount of NADPH is generated as well (Vander Heiden et al., 2009). Hence high glucose and high NAPDH levels are commonly found in cancer cells. Interestingly, in this study, we found that the accumulation of intracellular glucose and NADPH in cancer cells facilitated the activation of the polyol pathway (Figure 1H).

To determine whether the glucose level affects the activation of the polyol pathway, we compared the expressions of AKR1B1 and SORD in cancer cells cultured under different glucose levels. We found that glucose availability regulated the polyol pathway. We selected four different types of cancer cells with varying fructose production capabilities for this study. Under high glucose (25 mM) concentrations, the cancer cells significantly increased their AKR1B1 expression levels compared to those grown in low glucose (2.5 mM) (Figure 1I). The response of SORD expression to different glucose concentrations varied in the four cancer cell lines (Figure S1H). This observation may also provide a mechanistic explanation for the much-increased cancer risk in the diabetic patient population(Ohkuma et al., 2018). Increased blood glucose activates the polyol pathway, not only causing tissue damage due to increased production of sorbitol but also providing damaged cells with additional nutrient support to facilitate their malignant transformation.

### Fructose metabolism enhances glycolysis in cancer cells

To investigate the role of fructose metabolism in cancer cells, we compared the AKR1B1 KO (the polyol pathway deficient) A549 cells with their wild-type counterparts (control) in the Seahorse Cell Glycolysis Stress test and Mito Stress test assays (Bononi et al., 2017a). The assays showed drastically compromised glycolysis in the AKR1B1 KO cells (Figure 2A) but no significant difference in mitochondrial functions (Figure 2B) between the two cell lines. We then examined metabolites involved in glycolysis and mitochondrial respiration. Targeted metabolomics profiling showed that the glycolytic metabolites, such as serine (Amelio et al., 2014), pyruvate, and lactate, were significantly decreased in AKR1B1 KO cells (Figure 2C-F). Meanwhile, metabolites related to aerobic respiration, such as succinate, fumarate, malate, and aspartate (Sullivan et al., 2015) were slightly decreased or not significantly altered (Figure 2G-J and Figure S2A). These results indicated that fructose metabolism promoted glycolysis, leading to enhanced amino-acid (Figure 2C, 2G, 2F and Figure S2B) and pyruvate (Figure 2D) production. Pyruvate was first provided to the TCA cycle in mitochondria (Figure 2H-J), and after TCA metabolism reached the saturation point, the excess amount of pyruvate was converted to lactate (Figure 2E). We observed similar metabolic phenotypes in AKR1B1 silenced CY790 cells (Figure S2C-K), the wild-type of which showed high fructose production capability close to that of A549 cells.

**Figure 2.**
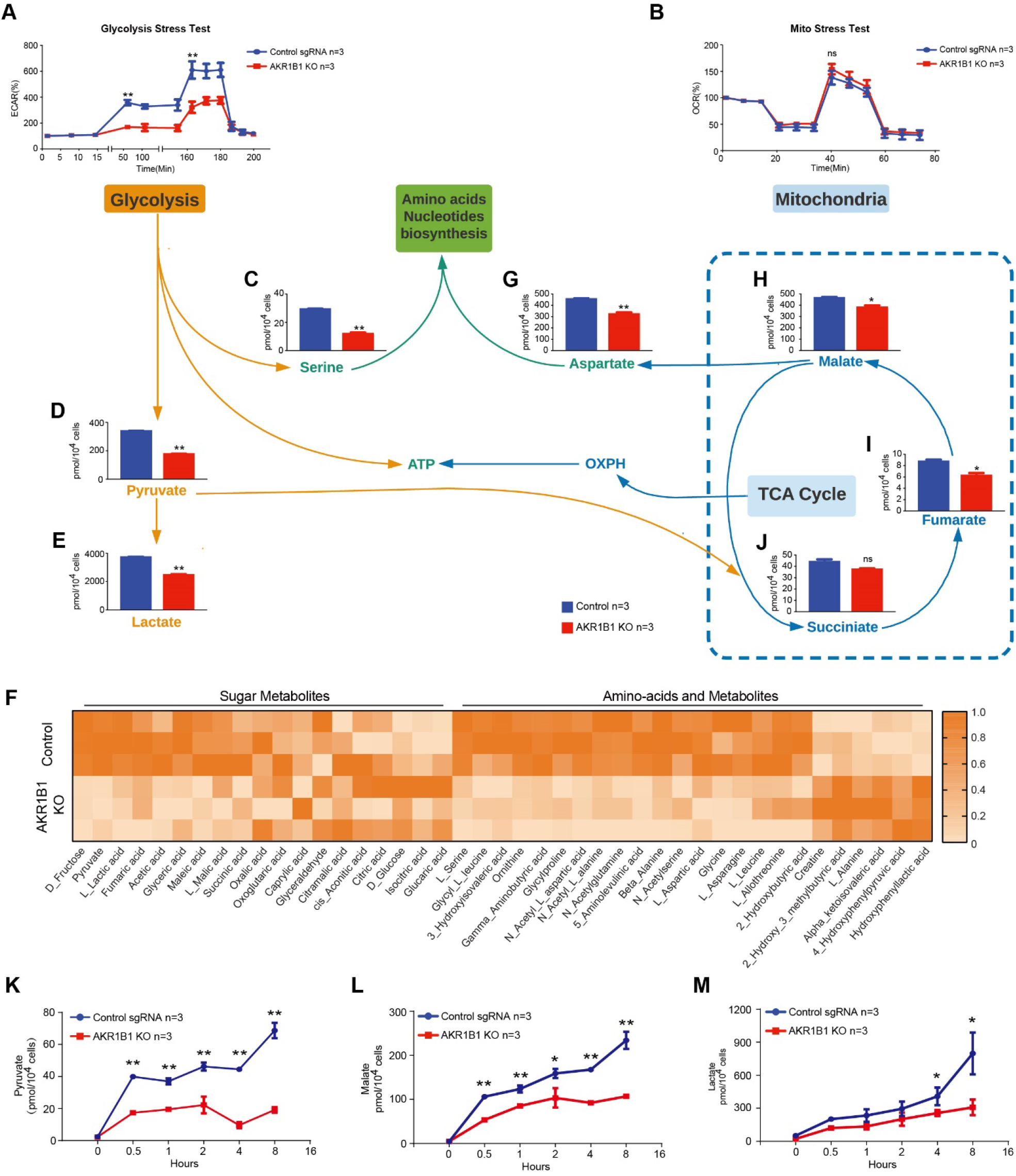
Fructose metabolism enhances glycolysis in cancer cells. **(A**) The Seahorse Cell Glycolysis Stress test showed compromised glycolysis in AKR1B1 KO A549 cells. **(B)** The Mito Stress test showed no significant change in mitochondrial function between A549 wild-type control and AKR1B1 KO cells. The direct metabolites from glycolysis, serine **(C)**, pyruvate **(D)**, and lactate **(E)** were significantly decreased in AKR1B1 KO cells. (**F**) Heat map of the results of metabolomics of wild-type control and AKR1B1 KO A549 cells. The metabolites are categorized into “sugar metabolites”, which includes metabolites related to glucose and fructose metabolism, and “amino-acid and metabolites”, which contains metabolites related to amino-acid metabolism. In comparison, TCA cycle-related metabolites, such as succinate **(J)**, fumarate **(I)**, malate **(H)**, and aspartate **(G)** were only slightly reduced in the KO cells. We also compared the flux of pyruvate **(K)**, malate **(L)**, and lactate **(M)** between wild-type control and AKR1B1 KO A549 cells when feeding with glucose after starving. The KO cells had slower rates in both of these metabolites. Data are represented as mean ± SEM. *: *t*-test p<0.05; **: *t*-test p<0.01.

Metabolic flux analysis showed that wild-type cells had a significantly faster pyruvate-producing rate than the polyol pathway-deficient cells (Figure 2K). The malate production curve showed a similar trend to pyruvate production (Figure 2L), suggesting that fructose metabolism accelerated pyruvate production and rapidly fueled the TCA cycle. The lactate production, however, increased at a much slower rate than those of pyruvate and malate in the wild-type cells (Figure 2M), suggesting that after the TCA cycle reached its saturation point, additional pyruvate derived from fructose metabolism was then converted to lactate. Such metabolic outcome is the phenomenon that was observed by Dr. Otto Warburg and perceived to be “aerobic glycolysis” of glucose metabolism in cancer cells about 100 years ago.

### The Warburg effect is a result caused by the synergy of glucose and fructose metabolism

To evaluate the contribution of fructose metabolism to the Warburg effect, we knocked out or silenced the Hexokinase-1 (HK1) and Hexokinase-2 (HK2) genes from A549 cells (Figures S3A and B). The hexokinases phosphorylate glucose to G-6-P (Figure 1H) and start the glycolysis. Both the hexokinases HK1 and HK2 are expressed in cancer cells. Knock out or silencing of HK1 and HK2 effectively blocked glucose metabolism in A549 cells (Xu et al., 2018) (Figures S3A-D). As described above, the A549 cells could process about 50% of glucose to fructose, i.e., the amounts of glucose and fructose are almost equal in this cell line. Blockage of glucose or fructose metabolisms in this cell line should thus be comparable.

Interestingly, the amount of ATP was drastically depleted in both glucose metabolism deficient and fructose metabolism defective cells at similar levels (Figure 3A and Figure S3E). We then compared the extracellular acidification rates (ECAR), a symbolic index of the Warburg effect, in the AKR1B1 KO and HK KO (or silenced) cells. The data showed that blockage of glucose or fructose metabolism resulted in a similar level of decrease in acidification in the A549 cancer cells (Figure 3B and Figure S3F), as well as pyruvate, lactate (Figure 3C and D), and other metabolites (Figure S3J-L). But the HK1/2 KO did not reduce the serine level in A549 cells as AKR1B1 KO (Figure 3E). It suggested that the serine in A549 cells might be mainly converted from fructose through D-glycerate (Duran et al., 1987).

**Figure 3.**
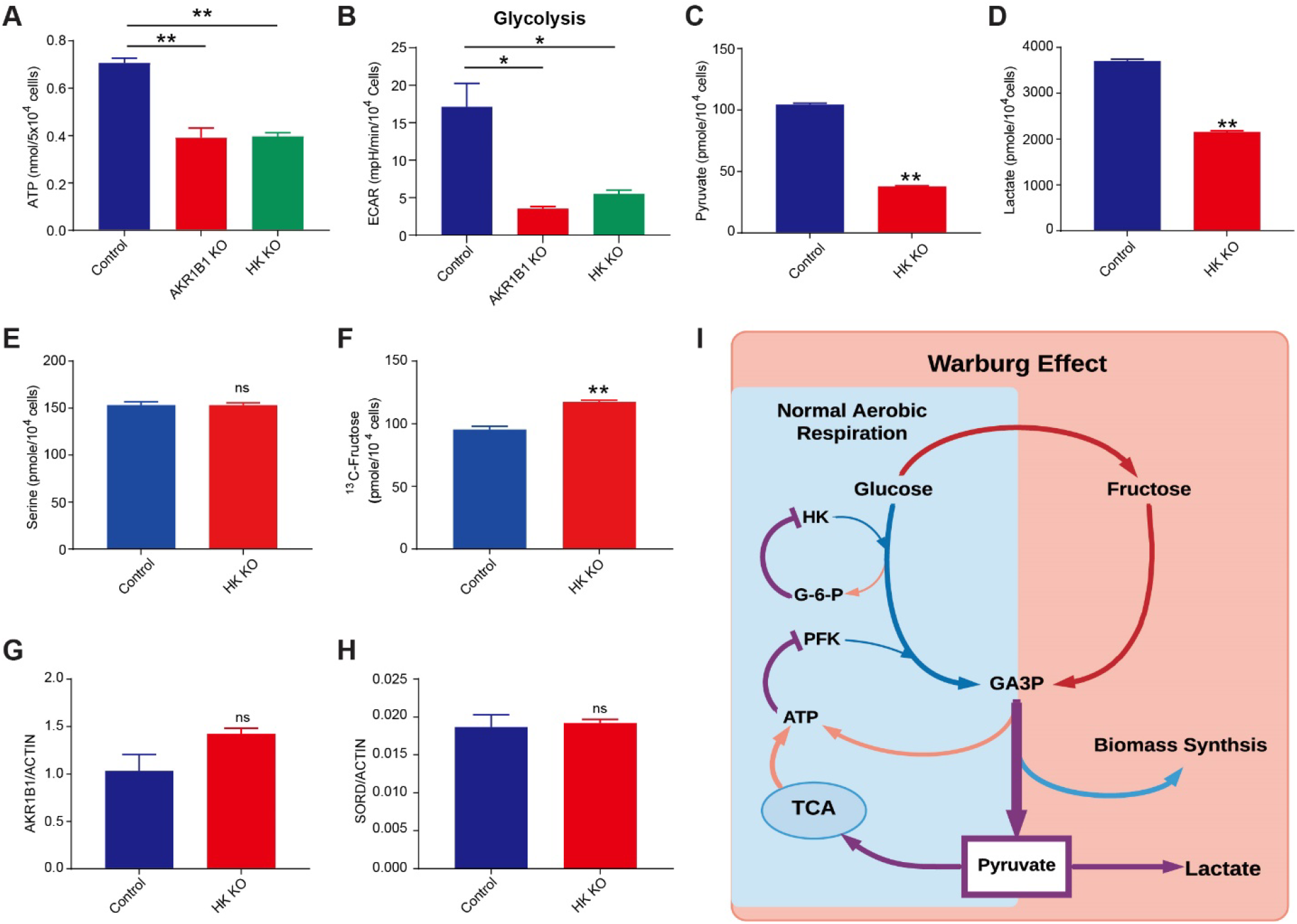
Fructose metabolism contributes to the Warburg effect. **(A**) Compromised fructose or glucose metabolism leads to diminished ATP production, as we observed in AKR1B1 KO cells as well as HK KO cells. We then compared glucose and fructose metabolisms’ effects on glycolysis using the Seahorse glycolysis stress test (**B**), as well as metabolomics as pyruvate (**C**), lactate (**D**), and serine (**E**). To evaluate how significant the glucose metabolism affects the polyol pathway, we tested fructose production in glucose metabolism compromised cells (**F)**, as well as expressions of AKR1B1 (**G**) and SORD (**H**). (**I**) Sketch of fructose metabolism contributions to the Warburg effect. Data are represented as mean ± SEM. *: *t*-test p<0.05; **: *t*-test p<0.01.

These results implied that glucose and fructose were equally capable of fueling aerobic glycolysis. As we know, glucose metabolism is a self-limited process with negative regulation of PFK and HK by glycolysis products such as ATP and G-6-P (Jeremy M Berg, 2002). To verify whether metabolites from glucose metabolism affected fructose conversion, we further tested the production of fructose and expressions of AKR1B1 and SORD in glucose metabolism deficient cells. The results showed that compared to control cells, neither fructose production nor AKR1B1 and SORD expressions drastically changed in HK KO cells (Figures 3F-H and Figures S3G, H). These implied that fructose metabolism was relatively independent of glucose metabolism.

As a tightly controlled process, glucose metabolism will reach a dynamic balance to supply a limited amount of materials and energy for regular cell metabolism. But in highly proliferating cancer cells, this is not sufficient. Under such circumstances, the fructose metabolism will bypass these control elements of classic glycolysis and keep providing glyceraldehyde 3-phosphate (GA3P) into the later glycolysis stage (Figure 3I). The additional GA3P from fructose metabolism will not predominantly fuel the TCA cycle, which is saturated through glucose metabolism. Instead, fructose metabolism will enhance glycolysis by increasing the biomass synthesis and production of lactic acid. In other words, the Warburg effect or aerobic glycolysis in cancer cells is achieved by a combined glucose and fructose metabolism.

### Cancer cells export endogenous fructose through the sugar transporter SLC2A8/GLUT8

Cancer cells absorb extracellular fructose predominantly through the fructose transporter, SLC2A5/GLUT5. Interestingly, we detected ^13^C-fructose in culture supernatants of all cell lines incubated with ^13^C-glucose, except CY790, a mesothelioma cell line (Figure 4A). This finding suggests that fructose produced in cancer cells can be transported to the extracellular environment. We compared the two mesothelioma cell lines, CY790 and CY289, one with a trace amount of fructose and the other with high concentration fructose (7 μM vs. 894 μM) found in the supernatant, respectively. The SLC2A5/GLUT5 and SLC2A8/GLUT8 expression levels were distinct between the two cell lines (Figure 4B). We detected SLC2A8 expression in CY289 but not in CY790 cells, suggesting that the difference in SLC2A8 expression between the two cell lines may be responsible for the different fructose concentrations in supernatants. We then knocked out the expression of SLC2A8 in A549 cells using a CRISPR/Cas9 technique (Figure 4C) as this cell line demonstrated strong fructose-exporting ability with the highest concentration of fructose in the culture supernatant among 14 cell lines (Figure 4A). The A549 SLC2A8 knockout cells showed significantly higher intracellular fructose levels and lower extracellular fructose levels than the A549 wild-type (Figure 4D), indicating that fructose export was effectively shut down. SLC2A8 is widely expressed in almost all tested cancer cell lines, as shown in Figure 4E, similar to the expression of the two genes encoding AKR1B1 and SORD in the polyol pathway. This finding was also validated by the TCGA human cancer database (Figure 4F), which reported that SLC2A8 was expressed across the spectrum of all types of human cancer.

**Figure 4.**
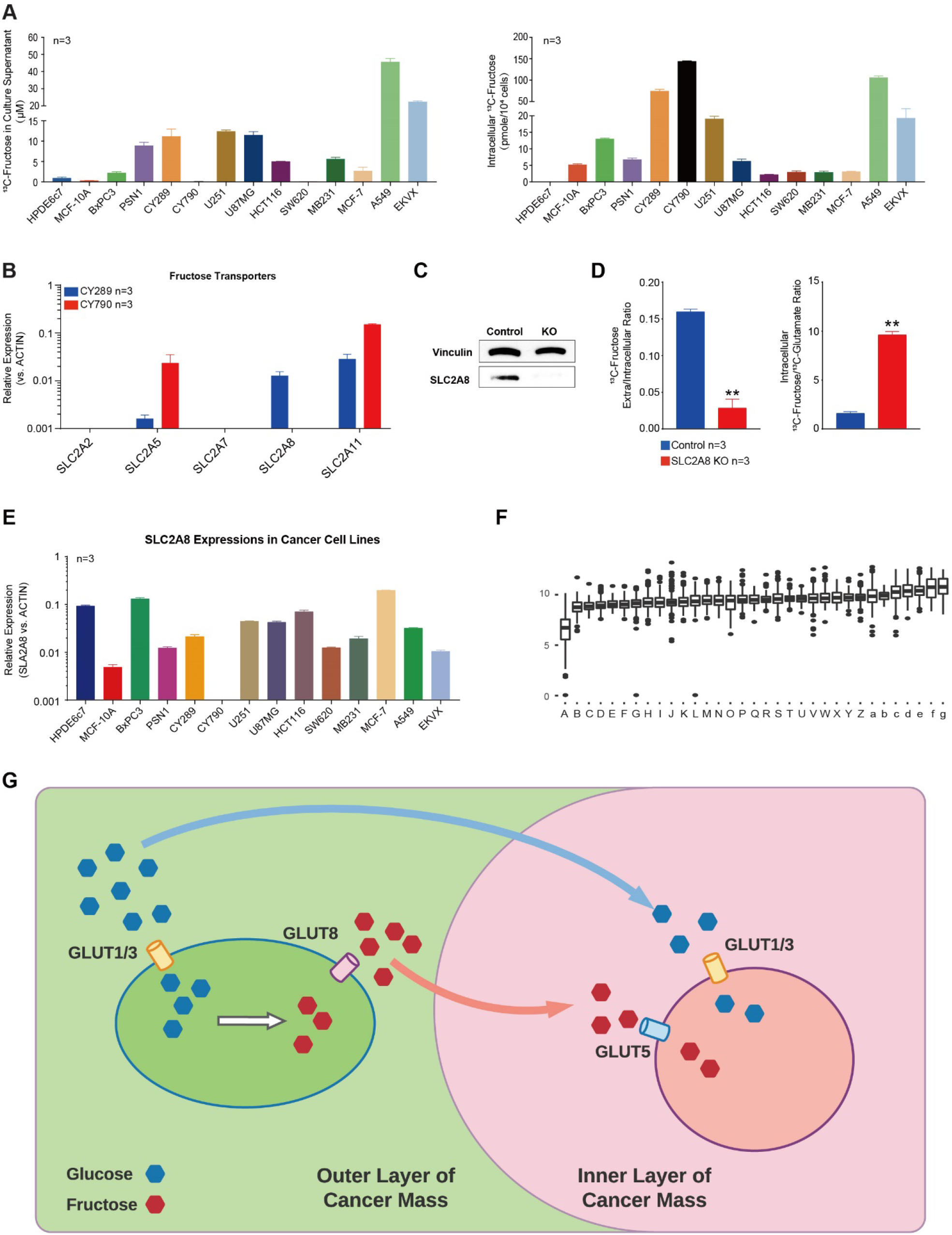
Cancer cells export endogenous fructose through the sugar transporter SLC2A8/GLUT8. **(A)** We found that cancer cells that produced fructose can export it to the extracellular environment, and that fructose distribution is varied in different cancer cells. Some cells tended to keep fructose in the cytoplasm, and others exported more fructose to the culture medium. This may be due to differences in fructose transporter expression in different cancer cells. **(B)** We tested potential fructose transporter(Debosch et al., 2014; Manolescu et al., 2005; Scheepers et al., 2005) expressions in two mesothelioma cell lines with different fructose distribution patterns and found that the expression of SLC2A8 was different between the two cell lines. **(C)** We generated SLC2A8 KO A549 cells by CRISPR/Cas9 technology and compared them with the control sgRNA group. **(D)** In SLC2A8 KO A549 cells, fructose export was highly suppressed, and the extracellular/intracellular fructose ratio was significantly reduced compared with the control group, while the intracellular fructose/glucose ratio was increased considerably. **(E)** SLC2A8 was also found to be widely expressed in cancer cells and (**F**) in human cancers according to the TCGA database, including: A. acute myeloid leukemia; B. thymoma; C. mesothelioma; D. thyroid carcinoma; E. cholangiocarcinoma; F. pancreatic adenocarcinoma; G. lung adenocarcinoma; H. breast invasive carcinoma; I. esophageal carcinoma; J. kidney clear cell carcinoma; K. head & neck squamous cell carcinoma; L. ovarian serous cystadenocarcinoma; M. lung squamous cell carcinoma; N. cervical & endocervical cancer; O. skin cutaneous melanoma; P. kidney papillary cell carcinoma; Q. uterine carcinosarcoma; R. pheochromocytoma & paraganglioma; S. stomach adenocarcinoma; T. brain lower grade glioma; U. glioblastoma multiforme; V. sarcoma; W. uterine corpus endometrioid carcinoma; X. diffuse large B−cell lymphoma; Y. testicular germ cell tumor; Z. prostate adenocarcinoma; a. bladder urothelial carcinoma; b. uveal melanoma; c. colon adenocarcinoma; d. rectum adenocarcinoma; e. liver hepatocellular carcinoma; f. kidney chromophobe; g. adrenocortical cancer. (**G**) Sketch of the fructose communication between cancer cells. Data are represented as mean ± SEM. **: *t*-test p<0.01.

This novel finding led us to hypothesize that fructose-producing cancer cells can make this additional energy source available to other cells in the tumor microenvironment. Generally, the nutrient supply is compromised inside a fast-growing tumor mass. However, with the polyol pathway and SLC2A8-mediated fructose exchange, it may be possible that the cancer cells in the outermost layer of the tumor can access circulating glucose and convert it to fructose and then export it into the tumor microenvironment for utilization by other cancer cells in the inner layer of the tumor cell mass (Figure 4G).

### Fructose enhances cancer cell proliferation and malignancy

To determine how fructose metabolism benefits cancer cells, we measured the cell responses to fructose in AKR1B1 KO cells and also cells overexpressed with AKR1B1. The AKR1B1 KO cells showed a decrease in malignant phenotypes *in vitro* and *in vivo*, while cells overexpressed in AKR1B1, on the other hand, demonstrated significantly increased rates of growth and tumor development in a mouse model (Figure 5 and Figure S4). To study the role of fructose metabolism via the polyol pathway in cancer cell growth, we generated AKR1B1 KO (Figure S1E) and AKR1B1 overexpressed cancer cell lines (Figures S4A) using CRISPR/Cas9 knockout and CRISPRa/Cas9-VPR overexpression technology. In the original AKR1B1 high expression cancer cells, such as A549 and CY790 cells, the CRISPRa/Cas9-VPR had a negligible effect on the AKR1B1 expression. In U87 and BxPC3 cells with moderate expression levels of AKR1B1, however, fructose production was increased by 3-5 fold with overexpression of AKR1B1 (Figure S4B).

**Figure 5.**
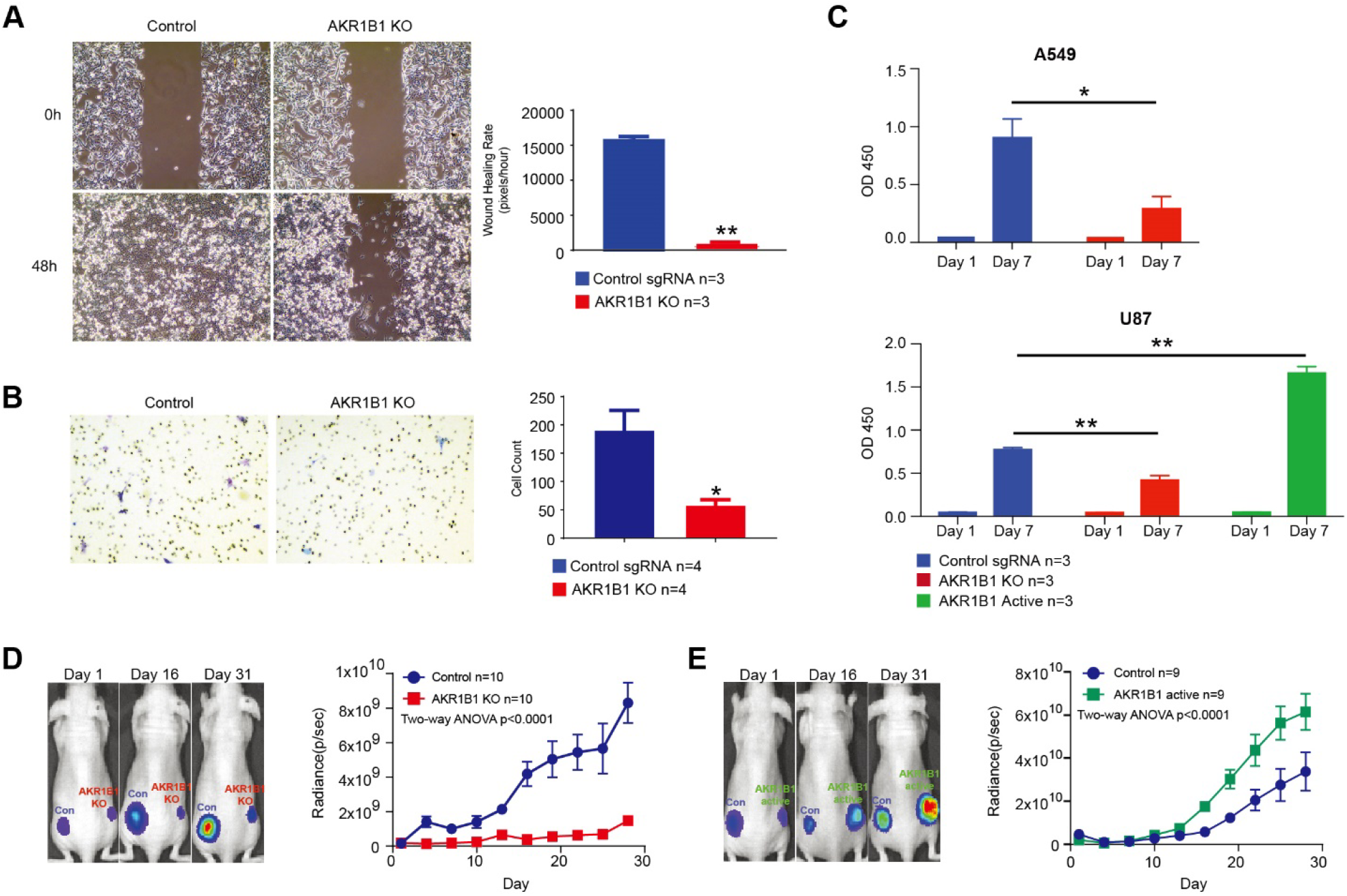
Fructose enhances cancer cell proliferation and malignancy. **(A)** The A549 AKR1B1 KO cells showed suppressed cell migration in the wound healing assay and **(B)** transwell assay. **(C)** The AKR1B1 KO cells slowed down cancer cell growths, and overexpression of AKR1B1 caused increased growth rates of U87 cells *in vitro*. **(D)** An *in vivo* xenograft model also showed growth suppression when AKR1B1 KO A549 cells were used, (**E)** there was increased cancer cell proliferation when U87 cells were overexpressed in AKR1B1. Data are represented as mean ± SEM. *: *t*-test p<0.05; **: *t*-test p<0.01.

We tested the migration and invasion ability of AKR1B1 KO A549 cells or AKR1B1 silenced CY790 cells. The knockout or silencing of AKR1B1 resulted in compromised cell motion in both the wound healing assay (Figure 5A and Figure S4C) and transwell assay (Figure 5B and Figure S4D), compared with wild-type control cells. *In vitro* cell proliferation tests showed that the growth rate of AKR1B1 KO or silenced cancer cells was highly suppressed (Figure 5C and Figures S4E, F). On the other hand, overexpression of AKR1B1 not only increased fructose production but also significantly increased tumor growth rate (right panels of Figure 5C and Figure S4E). We then investigated the effect of the polyol pathway on cancer proliferation *in vivo*. We injected 0.1 ×10^6^ wild-type or gene manipulated (AKR1B1 KO, overexpression, or AKR1B1 silenced) cancer cells, which constitutively expressed luciferase, in nude mice on day 0, respectively. Luciferase activity of each tumor increased with tumor mass. Every three days, we measured the luminescence intensity of tumors by D-luciferin injection. Compared with wild-type cells, AKR1B1 compromised (KO or silenced) cancer cells showed significantly reduced tumor growth rates (Figure 5D and Figures S4G-I) while AKR1B1 overexpression promoted cancer cell growth (Figure 5E, and Figure S4J). These results confirmed that fructose metabolism enhanced cancer malignancy.

## DISCUSSION

Our study identified the intra- and extracellular polyol pathways as a crucial fructose-producing mechanism that is important for cancer-mediated metabolic reprogramming leading to enhanced glycolysis, cell proliferation, and malignancy. Compared with glucose, fructose metabolism results in accelerated glycolysis and an increased ATP amount, demonstrating a typical phenotype of increased glycolytic flux. We suspect that cancer metabolism involving the intra-/extracellular polyol pathway-derived fructose may constitute an essential part of the Warburg effect. Under the aerobic condition, the abundant ATP from the TCA cycle limits the glycolytic rate. Almost all of the glycolysis production will be exhausted by the TCA cycle with nothing remaining to convert to significant amounts of lactate. In cancers, uptake of large amounts of glucose through overexpressed GLUT1 and/or GLUT3 activates the polyol pathway and fructose metabolism, overcoming glycolysis limitations and producing an excess amount of glycolytic metabolites for biomass and lactate production. It results in the acidification of cancer cells under aerobic conditions. Additionally, cancer cells have a selective advantage to utilize fructose to support their rapid proliferation because fructose lacks cellular metabolic homeostasis machinery, unlike glucose, which is strictly regulated by either insulin or glucagon. As the polyol pathway is not cancer-specific (comparable levels of AKR1B1 and SORD were found in normal epithelial cell lines, Figures S1A, B), we hypothesize that it profoundly impacts all fast-growing cells, including activated immune cells, germinal cells, and stem cells.

We also found in this study that high glucose levels stimulated the expression of AKR1B1. Therefore, it is not difficult to understand that the polyol pathway is activated to lower the blood glucose levels in diabetic patients and, thus, implicated in diabetic complications. The intermediate molecule in the polyol pathway, sorbitol, cannot cross cell membranes and, when it accumulates from increased cellular production, causes microvascular damage to the retina, kidney, and nerves (Yabe-Nishimura, 1998). Given the recent findings that increased fructose level promotes cancer malignancy, the polyol pathway’s activation may be responsible for the increased risk of cancer development in the diabetic population.

As the polyol pathway is not essential to energy metabolism in normal cells via the glycolysis pathway, inhibition of AKR1B1 activity may represent a novel, effective, and safe strategy for cancer therapy and preventive cancer treatments for diabetic patients. Currently, several AKR1B1 inhibitors are undergoing clinical trials to treat diabetic complications(Goto et al., 1995; Quattrini and La Motta, 2019). These compounds may potentially be useful as direct treatments for cancer and preventive cancer treatments for diabetic patients.

To summarize, the cancer cells utilize the polyol pathway to produce fructose and gain more energy and biomass from fructose metabolism than the equivalent amount of glucose. Fructose metabolism in cancer cells causes an increased amount of lactate and metabolic intermediates for biomass, demonstrating a typical glycolytic phenotype and providing an essential contribution to the Warburg effect.

## Methods

### Cell Culture

A549 (lung carcinoma), EKVX (lung adenocarcinoma), SW-620 (colorectal adenocarcinoma), HCT116 (colorectal carcinoma), U251 (malignant glioblastoma), MCF7 (breast adenocarcinoma), and MDA-MB-231 (breast adenocarcinoma) cells were obtained from NCI-60 cell storage of the University of Hawaii Cancer Center. BxPC3 (pancreas adenocarcinoma), PSN1 (pancreas adenocarcinoma), and U87 MG (malignant glioblastoma) cells were ordered from Sigma-Aldrich. HPDE6c7 (pancreatic duct epithelial cell), and MCF-10A (breast fibrocystic disease) cells were ordered from American Type Culture Collection (ATCC). CY289 and CY790 (mesothelioma) cells were gifts from Dr. Carbrone and Dr. Yang’s lab. All cell lines were adapted to culture and maintained in DMEM (Thermo Fisher Scientific) with 10% FBS under 37°C and 5% CO_2_.

### Gene Knock Out and Activation by CRISPR/Cas9

We used Dharmacon Edit-R All-in-one Lentiviral and Edit-R CRISPRa Lentiviral systems to create gene knock out (KO) and gene overexpression cell lines. The sgRNA sequences are as follows: CGACCTGAAGCTGGACTACC (GSGH11935-247543511, for Human AKR1B1 KO, Dharmacon), GCCGCGGCGGCCTTCCCC AA (GSGH11887-247076986, for Human AKR1B1 overexpression, Dharmacon), GACAACTATACCTGCCAGGT (GSGH11935-247493945, for Human SLC2A8 KO, Dharmacon), GTTCAAGATGGCGCCCAGTG (GSGH11935-247518767, for Human HK1, Dharmacon), TAAGCGGTTCCGCAAGGAGA (CS-HCP001744-LvSG03-1-B, for Human HK2, GeneCopoeia). The shRNA sequences are as follows: GGTGGAGATGATCTTAAACAA (HSH005861-LVRU6GP, for Human AKR1B1, GeneCopoeia), CAATGCCTGCTACATGGAGGA (CS-HSH1530L-LVRU6GP-01, for Human HKs, GeneCopoeia). Lentivirus was packed using the Dharmacon Trans-Lentiviral ORF Packaging Kit with Calcium Phosphate (TLP5916), and cells were infected to create stable cell lines according to the manufacturer’s instruction.

### Western Blotting

Immunoblotting was performed as previously described (Chen et al., 2016). Briefly, the transfer-ready membrane was blocked 1 hour in TBS-T containing 5% nonfat milk at room temperature, followed by incubation with primary antibody at 4°C overnight. The anti-AKR1B1 antibody (Abcam, ab62795), anti-SORD antibody (Abcam, ab185705), anti-SLC2A8 antibody (LifeSpan BioSciences, LS-C757596), and anti-Vinculin antibody (Cell Signaling Technology, E1E9V) were used at 1:1000 dilution respectively. The secondary antibody was horseradish peroxidase-conjugated anti-rabbit antibody used at a 1:5,000 dilution. Vinculin was used as a protein loading control.

### RT-PCR

RNA extraction and reverse transcription were performed by TRIzol (Thermo Fisher Scientific) and iScript™ cDNA Synthesis Kit (Bio-Rad Laboratories) according to the manufacturer’s instructions. PCR primer sequences were as follows. Primers for AKR1B1 PCR: TACTCAGCTACAACAGGAACTG, AGGCAAGAAACACAG GTATAGG, probe: TTGTTGAGCTGTACCTCCCACAAGG; Primers for SORD PCR: ACTCCAGAGCCAAAAGAGC, CATCCTCAGCAAGACCTCAT, probe: AGGATAGTTCTCCAGGCGCAAGTC; Primers for SLC2A8 PCR: GCCAAG TTCAAGGACAGCA, ATGACCACACCTGACAAGAC, probe: ATGATGAGA GCCGCCACAGCT; Primers for ACTB PCR: GCGAGAAGATGACCCAGAT, CCAGTGGTACGGCCAGA, probe: CCATGTACGTTGCTATCCAGGCTGT; Primers for SLC2A5 PCR: CAAGCATGGAGCAACAGGAT, GAAGGATGA CCCAAAGGCA, probe: AGCATGAAGGAAGGGAGGCTGAC; Primers for SLC2A2 PCR: CTGGAAGAAGCATATCAGGACT, GCTGATGAAAAGTGC CAAGTG, probe: CTGAGAGCGGTTGGAGCAATTTCAC; Primers for SLC2A7 PCR: CCGTCTCCATGTTTCCTCTG, CCACTTTGCTGACTCCCATC, probe: CCCACGAGCAATGACCCCAACA; Primers for SLC2A11 PCR: GTCAGCAGC AATCCTGTTTG, GTACATGGGCTGGATGTTCA, probe: AGCAGTCTTCCC AGCATGATCATCTC. We performed real-time PCR with the iTaq™ Universal Probes Supermix (Bio-Rad Laboratories) by Lightcycle 480 (Roche Molecular Systems).

### Seahorse Extracellular Flux Assays

Gene-modified cancer cells were maintained in serum-free HuMEC Ready Medium before the assay. The Seahorse XF Glycolysis Stress Test Kit and Cell Mito Stress Test Kit (Agilent) were used to measuring cell glycolysis and mitochondrial function, respectively, using the Agilent Seahorse XF96 analyzer(Bononi et al., 2017a) following the manufacturer’s instructions.

### Measurements of AKR1B1 and SORD in human serum

Peripheral blood specimens from 10 healthy donors, 16 lung cancer, 8 colorectal cancer, and 22 multiple myeloma patients were obtained with consent at the Second Xiangya Hospital of Central South University, Blood Diseases Hospital of Chinese Academy of Medical Science & Peking Union Medical College, Hunan Cancer Hospital, Hunan Provincial People’s Hospital and the Second People’s Hospital of Hunan Province during the period 2016-2020. Serum was prepared from peripheral blood. The concentrations of AKR1B1 and SORD in serum were measured using Human Aldose Reductase (AR) ELISA Kit (Catalog No. ml024614, Mlbio, Shanghai, China) and Human Sorbitol Dehydrogenase (SORD) ELISA Kit (KL-SORD-Hu, KALANG, Shanghai, China) according to their manufactory instructions, respectively.

### Cell Proliferation Assay

All gene-modified cancer cells were first adapted to HuMEC Ready Medium (Thermo Fisher Scientific) and then plated at 1000 cells/well in HuMEC Ready Medium followed by the culture at 37°C under 5% CO_2_. A cell proliferation assay was performed by using the Cell Counting Kit-8 (CCK-8) (Dojindo Molecular Technologies) after the indicated time point of culture. 10 μL of the CCK8 dye was added to each plate well and then incubated for 1 hour at 37°C under 5% CO_2_. The optical density (OD) was then read at 450 nm directly using a microplate reader.

### *In vitro* Cell Migration and Invasion Assays

The ibidi Culture-Insert 4 Well in µ-Dish (ibidi, 80466) was used for the wound healing assay, and the Corning BioCoat Matrigel Invasion Chamber (Corning, 354480) was used for the transwell assay according to the manufacturer’s instructions. Gene-modified cancer cells were seeded in HuMEC Ready Medium and cultured at 37°C under 5% CO_2_. The images of wound healing data were analyzed by NIH ImageJ.

### Xenograft Formation

Nude mice (4-6 weeks old) from Charles River Laboratory were injected right and left sides respectively with 200 μL Matrigel (Corning) slurry (prepared at a 1:1 ratio with 1X PBS) containing 0.1 million control or gene manipulated cancer cells that constitutively expressed luciferase (AMS Biotechnology Limited). Luciferase activity of each tumor increased with tumor mass by the time. Every three days, we measured the luminance intensity of tumors by intraperitoneal injection of 50 mg/mouse D-luciferin (Thermo Fisher Scientific) into anesthetized mice, followed by the detection of live images using the Xenogen IVIS (PerkinElmer) at 30 min post-injection.

### Analysis of TCGA data

The Cancer Genome Atlas (TCGA) data were downloaded from the UCSC Xena platform [https://doi.org/10.1038/s41587-020-0546-8]. Boxplots were plotted using ggplot2 3.2.1 under R 3.6.1. Kaplan-Meier survival analyses were performed by UCSC Xena [https://ucsc-xena.gitbook.io/project/overview-of-features/kaplan-meier-plots].

### In vitro Glucose-Sorbitol conversion

20ng/mL of BSA control protein, recombinant human AKR1B1(Novus Biologicals, LLC, NBC1-22925) or AKR1C1(R&D Systems, 6529-DH-020) was added into a reaction system composed of 10μM NADPH (Roche, 10107824001) and 25mM ^13^C-glucose (Cambridge Isotope Laboratories) in PBS. The reaction occurred at 37°C for 24 hours before harvest. The ^13^C-sorbitol in each reaction was quantified by UPLC-QTOFMS (Dong et al., 2016).

### Chemicals and reagents

Diisopropylethylamine (DIPEA), 3-nitrophenylhydrazine (3-NPH)·HCl, 1-ethyl-3-(3-dimethylaminopropyl) carbodiimide (EDC) HCl, and pyridine (HPLC grade), glucose, fructose, glucose 6-phosphate, fructose 6-phosphate, glucose 1-phosphate, fructose 1-phosphate, fructose 1,6-bisphosphate, glyceraldehyde 3-phosphate, dihydroxyacetone phosphate, 2-phosphoglycerate, pyruvate, 6-phosphogluconate, ribulose 5-phosphate, ribose 5-phosphate, sedoheptulose 7-phosphate, erythrose 4-phosphate, citrate, isocitrate, succinate, fumarate, malate, lactate, α-ketoglutarate, oxaloacetate, serine, glutamate, glutamine, aspartate, hydropyruvate were purchased from Sigma-Aldrich (St Louis, MO). Optima LC-MS grade water, methanol (MeOH), isopropanol, formic acid and acetonitrile (ACN) were purchased from Thermo Fisher Scientific (Waltham, MA). [13C6]-glucose and [13C6]-fructose D-Glucose U-13C6 and D-Fructose U-13C6 were purchased from Cambridge Isotope Laboratories (Tewksbury, MA). FBS: Premium Grade Fetal Bovine Serum (VWR, 97068-085), Performance Plus Fe tal Bovine Serum (gibco, 16000-036), Charcoal Stripped Fetal Bovine Serum (gibco, A33821-01), Dialyzed Fetal Bovine Serum (gbico, a33820-01). Heat Inactive was performed under 56 °C for 30 min. DMEM (gibco, 11966-025). Epalrestat (Cayman Chemical, 15214). 13C-Glucose (Cambridge Isotope Laboratories, CLM-1396). Amino Acids: L-Arginine, Hydrochloride (MilliporeSigma, 181003), Histidine hydrochloride (MilliporeSigma, H0755000), L-Lysine (MilliporeSigma, 62840), L-Phenylalanine (MilliporeSigma, P5482), L-Valine (MilliporeSigma, V0513). PBS (gibco, 14190-144).

### Standard solutions

Each standard compound was accurately weighed and dissolved in water to prepare stock solutions with a final concentration of 20 mM for each standard. The final working standard solution mix contained 28 metabolites (detailed in Table 1) with a final concentration of 400 μM/metabolite. For quantification, a 7-point calibration curve was constructed by using a series of 1:2 serial dilutions from the highest concentration of calibration mixture with water. The stock solutions were kept at −20 °C.

### Stable isotope tracer analysis and metabolite quantification

Cell culture, labeling, and sample collection: All tested cells were cultured in DMEM containing 10% FBS under 37°C/ 5% CO_2_ before treatment. After washing with PBS, all tested cells were treated with DMEM containing 25mM D-Glucose U-13C6 (Cambridge Isotope Laboratories (Tewksbury, MA))and 2% BSA under 37°C/5% CO_2_ for 24 hours or indicated time points for flux rate tests. ^13^C labeled metabolites in medium and cell pellets were identified and quantified by using UPLC-QTOF-MS.

UPLC-QTOF-MS analysis: Cells were washed twice with phosphate-buffered saline (Thermo Fisher Scientific) (pH 7.4) and then carefully scraped into 1.5 mL safe lock centrifuge tubes with 10 mg of beads (0.5 mm). Metabolites were extracted by adding 0.5 mL of 50% MeOH (−20°C), followed by homogenization for 3 min using a Bullet Blender Tissue Homogenizer (Next Advance, Inc., Averill Park, NY) and centrifugation at 13, 500 g for 10 min at 4°C.

The resulting 50 μL of supernatant and also cell culture medium samples were transferred to 1.5 mL tube and mixed with 10 μL of 200 mM 3-NPH solution and 10 μL of the mixed 96 mM of EDC/pyridine methanolic solution. Derivatization was conducted by incubation at 30°C for 1 h before evaporation to dryness under nitrogen. 400 μL of 50% aqueous methanol was used to re-suspend the samples. The supernatants were used for UPLC-QTOF-MS analysis according to previous reports with minor modifications(Bononi et al., 2017b; Paglia et al., 2012).

### Cell Metabolomics Analysis

Cell culture and sample collection: Cells were adapted to HuMEC Ready Medium and fed by fresh medium 24h before harvest for metabolomics assay. To perform the metabolite flux test, cells were starved for 16 hours in DMEM containing 2% BSA without glucose before feeding them with the HuMEC Ready Medium. Cells were then harvested cells at indicated time points.

Cell Metabolite extraction and derivatization: Cells were washed twice with phosphate-buffered saline (Mediatech, Manassas, VA) (1 mM pH 7.4) and then carefully scraped into 1.5 mL safe lock centrifuge tubes with containing 10 mg of beads (0.5 mm). Metabolites were extracted by adding 0.5 mL of 50% MeOH (−20°C), followed by homogenization for 3 min using a Bullet Blender Tissue Homogenizer (Next Advance, Inc., Averill Park, NY) and centrifugation at 13, 500 g for 10 min at 4°C. Derivatization was performed according to the method of Han et al.(Han et al., 2016) with minor modifications. 50 μL of supernatant or standard was transferred to a 1.5 mL tube and mixed with 10 μL of 200 mM 3-NPH solution, and 10 μL of the mixed 96 mM of EDC/pyridine methanolic solution. Derivatization was conducted by incubation at 30°C for 1 h before evaporation to dryness under nitrogen. 400 μL of 50% aqueous methanol was used to re-suspend the samples. The derivatives were analyzed via UPLC-QTOF-MS.

Cell extracts analysis by ultra-performance liquid chromatography/xevo G2 quadrupole time-of-flight tandem mass spectrometry (Figure S5): The metabolites in culture medium and cell pellets were analyzed using the ACQUITY UPLC I-Class System coupled with a Xevo G2-S QTOF (ACQUITY UPLC-Xevo G2 QTOF, Waters Corp., Milford, MA). Chromatographic separation of phosphate and carboxylate was performed on the ACQUITY UPLC with a conditioned autosampler at 10°C, using an Acquity BEH C18 column (100 mm × 2.1 mm i.d.,1.7 μm particle size) (Waters, Milford, MA, USA). The column temperature was maintained at 40°C. The mobile phase consisting of water with 5 mM DIPEA in water (solvent A) and acetonitrile/isopropanol=7/3 (solvent B) was pumped at a flow rate of 0.3 mL/ min. The gradient elution program was as follows: 0–8 min, 1-20% B; 8–16 min, 20–98% B; 16.0–16.1 min, 98–1% B; 16.1–18.0 min, 1% B for equilibration of the column. The injection volume was 5 μL. The Xevo G2 QTOF mass spectrometer was used in negative ESI mode for data acquisition using UPLC/MS. Typical source conditions were as follows: capillary voltage, 3.0 kV; sample cone, 40 V; source temperature, 120 °C; desolvation temperature 450 °C; cone gas flow rate 50 L/h; desolvation gas (N2) flow rate 900 L/h. All analyses were performed using the lockspray, which ensured accuracy and reproducibility. Leucine–enkephalin (5 ng /mL) was used as the lockmass generating a reference ion in positive mode at m/z 554.2615 and was introduced by a lockspray at 10 μL/min for accurate mass acquisition. Data acquisition was achieved using MS. The injection volume was 5 μL.

As shown in Extended Data Figure 5, a total of 28 phosphate and carboxylate metabolites were detected by UPLC-QTOF-MS. Extended Data Table 1 showed the identification of metabolites in cells using UPLC-QTOF-MS.

### Statistics and reproducibility

Statistical analyses were performed using GraphPad PRISM version 8.0 software. A two-tailed t-test assessed the statistical significance of data unless noted otherwise, and the p > 0.05 was considered not significant (ns). Data represent mean ± sem as indicated in figure legends. All experiments for which data are displayed as a dot plot, all values from each repeat of the experiment are displayed together. For all experiments shown, similar results were obtained from at least three biologically independent experiments.

## AUTHOR CONTRIBUTIONS

W.J. conceptualized the study. W.J. and B.H. designed the studies. B.H., M.W. and J.Z. performed and analyzed biological experiments. B.H., G.X. and L.W. performed and analyzed metabolomics experiments. W.J. and B.H. wrote the manuscript. W. C., C. R., W.Z., J.Z. and H.Y. edited the manuscript.

---

**Figure S1.**
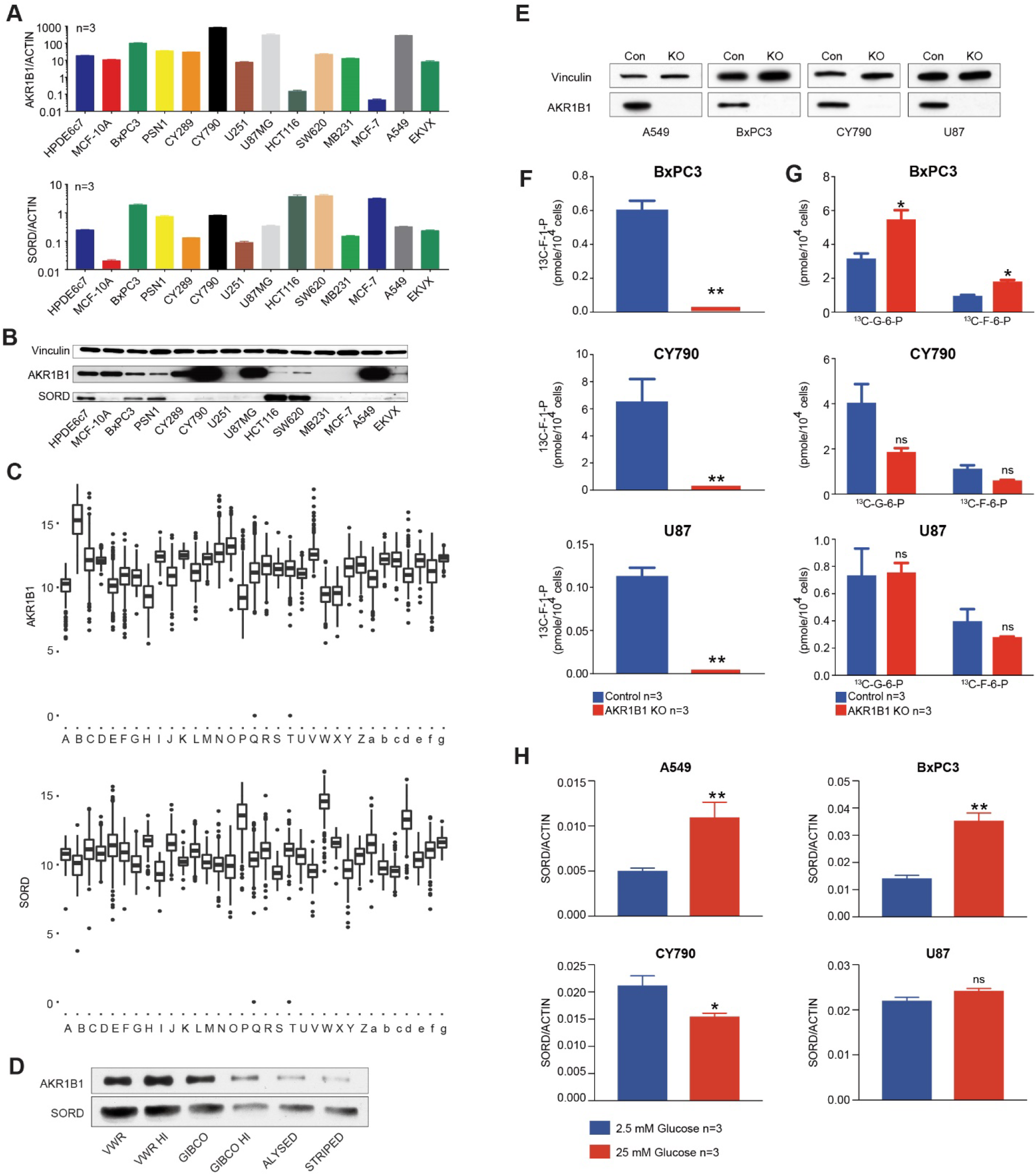
**(A-C)** Key enzymes of the polyol pathway, AKR1B1 and SORD, are widely expressed in multiple cancer cell lines at both mRNA **(A)** and protein **(B)** levels. **(C)** A TCGA database also showed expressions of these two genes in multiple human cancers. The cancer types are listed below: A. acute myeloid leukemia, B. adrenocortical cancer, C. bladder urothelial carcinoma, D. brain lower grade glioma, E. breast invasive carcinoma, F. cervical & endocervical cancer, G. cholangiocarcinoma, H. colon adenocarcinoma, I. diffuse large B-cell lymphoma, J. esophageal carcinoma, K. glioblastoma multiforme, L. head & neck squamous cell carcinoma, M. kidney chromophobe, N. kidney clear cell carcinoma, O. kidney papillary cell carcinoma, P. liver hepatocellular carcinoma, Q. lung adenocarcinoma, R. lung squamous cell carcinoma, S. mesothelioma, T. ovarian serous cystadenocarcinoma, U. pancreatic adenocarcinoma, V. pheochromocytoma & paraganglioma, W. prostate adenocarcinoma, X. rectum adenocarcinoma, Y. sarcoma, Z. skin cutaneous melanoma, a. stomach adenocarcinoma, b. testicular germ cell tumor, c. thymoma, d. thyroid carcinoma, e. uterine carcinosarcoma, f. uterine corpus endometrioid carcinoma, g. uveal melanoma. **(D)** AKR1B1 and SORD proteins in different FBSs. **(E)** CRISPR/Cas9 technology was used to successfully knocked out the AKR1B1 gene from multiple cancer cells as shown by Western blot, which indicated no detectable AKR1B1 in KO cancer cells. **(F)** Fructose specific metabolite, fructose-1-phosphate (F-1-P), highly depended on the polyol pathway. Only a trace amount of F-1-P can be detected in AKR1B1 KO cancer cells. **(G)** The compromised polyol pathway did not significantly affect glucose metabolism in most tested cancer cells. The amounts of glucose specific metabolites, glucose-6-phosphate and fructose-6-phosphate (G-6-P and F-6-P), only slightly increased in BxPC3 and had no significant change in other tested AKR1B1 KO cancer cells. We tested the regulation of expressions SORD **(H)** by glucose level in different cancer cells. *: *t*-test p<0.05; **: *t*-test p<0.01.

**Figure S2.**
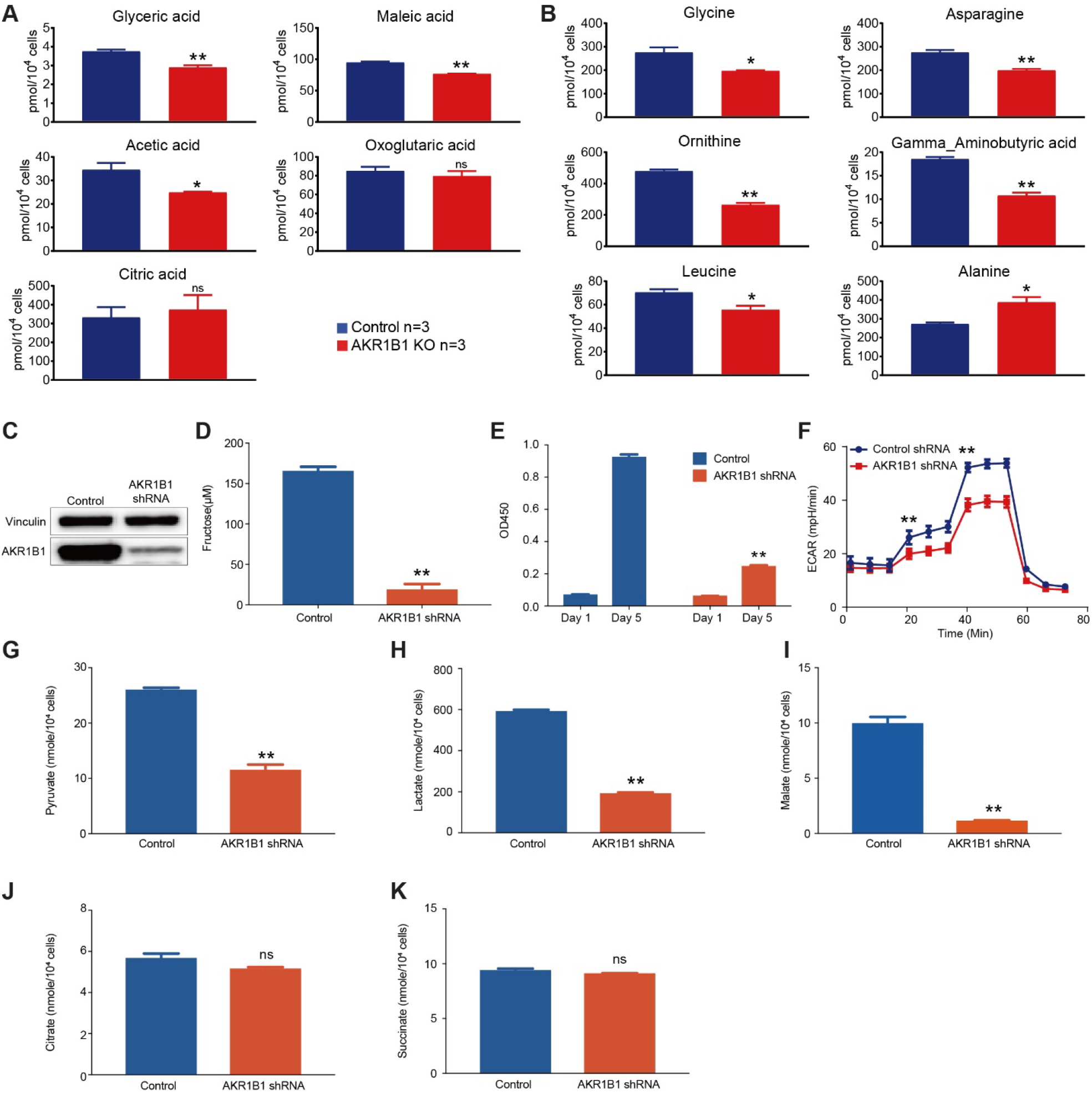
(**A**) Metabolites related to glucose and fructose metabolism. (**B**) Metabolites related to amino-acid metabolism. **(C)** We silenced the AKR1B1 gene expression from CY790 cells, as shown by Western blot, which indicated reduced AKR1B1 in shRNA treated cells. **(D)** AKR1B1 silencing reduced fructose production in CY790 cells. **(E)** The AKR1B1 silencing slowed down CY790 cell growth *in vitro*. **(F**) Seahorse Cell Glycolysis Stress test showed compromised glycolysis in AKR1B1 silenced CY790 cells. The pyruvate **(G)**, lactate **(H)**, and malate **(I)** were decreased in AKR1B1 silenced cells. While citrate **(J)** and succinate **(K)** were not significantly reduced in the AKR1B1 silenced cells. *: *t*-test p<0.05; **: *t*-test p<0.01.

**Figure S3.**
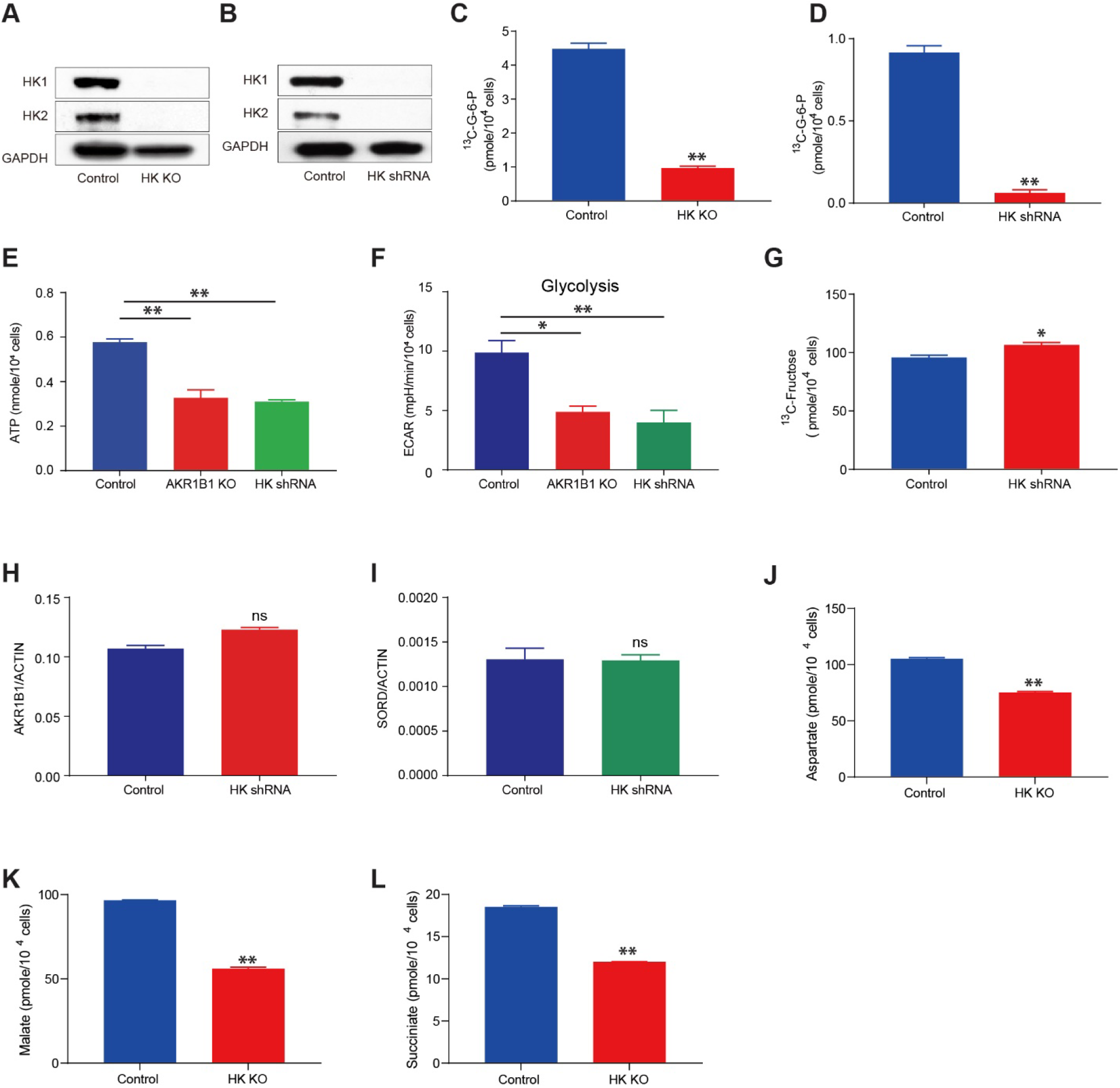
(**A** and **B**) HK1 and HK2 expression in HK KO and silencing A549 cells. (**C** and **D**) G6P, the downstream metabolite of glucose in glycolysis, is highly reduced in HK KO and silencing A549 cells. Compromised ATP production **(E)** and ECAR (**F**) were observed in HK silencing cells. (**G**) Fructose production was not significantly changed in HK silencing A549 cells. (**H** and **I)** AKR1B1 and SORD expressions in HK silencing A549 cells. Aspartate (**J**), Malate (**K**) and Succiniate (**L**) in A549 control and HK KO cells. *: *t*-test p<0.05; **: *t*-test p<0.01.

**Figure S4.**
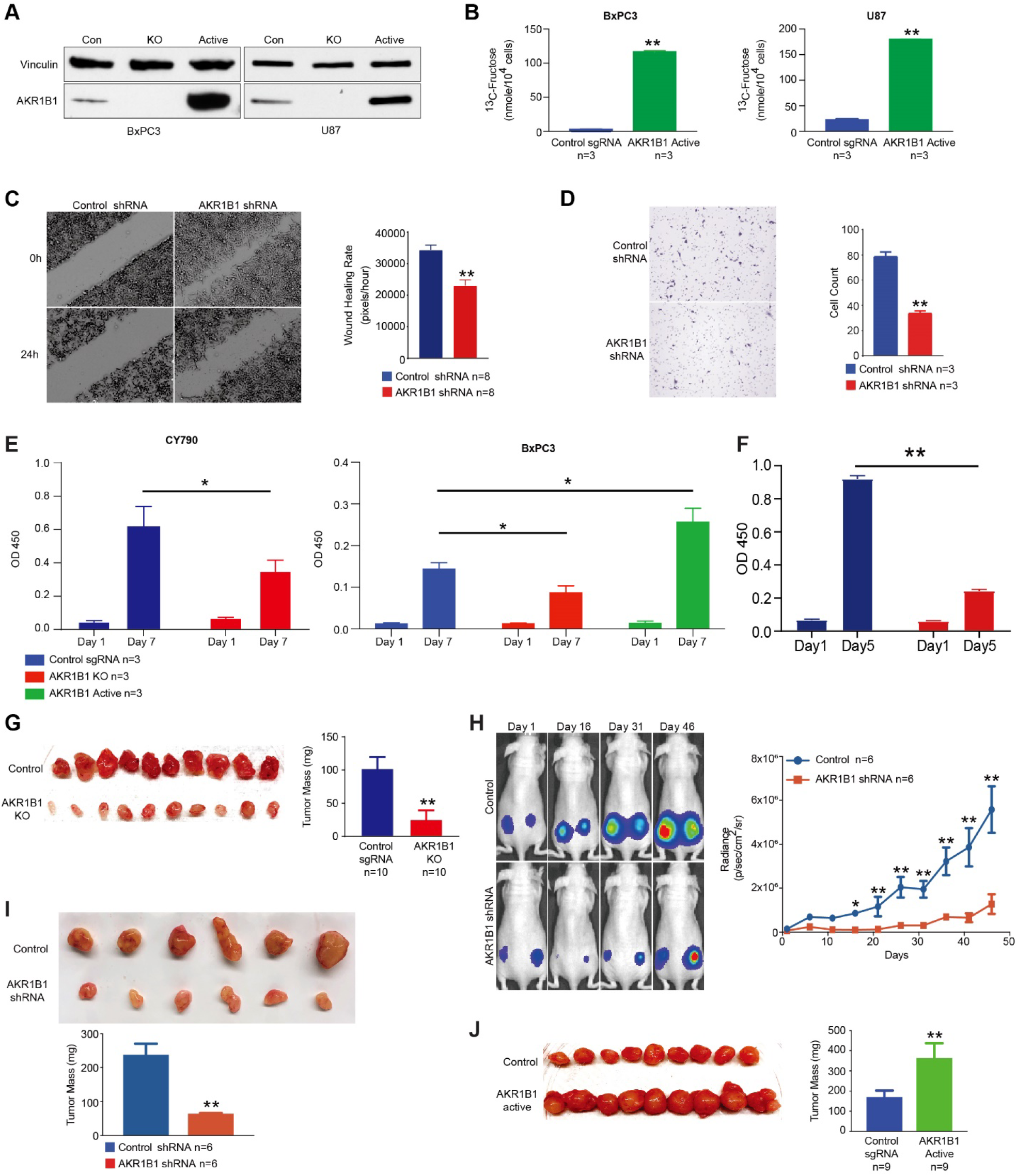
(**A**) CRISPRa/Cas9-VPR technology was used to overexpress AKR1B1 in cancer cells, and it significantly increased AKR1B1 expression relative to controls. **(B)** Overexpression of AKR1B1 increased fructose productions in U87 and BxPC3 cells. **(C)** The CY790 AKR1B1 silenced cells showed suppressed cell migration in the wound healing assay and **(D)** transwell assay. **(E)** The AKR1B1 KO cells slowed down cancer cell growths, and overexpression of AKR1B1 caused increased growth rates of BxPC3 cells *in vitro*. **(F)** The AKR1B1 silenced CY790 cells slowed down cancer cell growths *in vitro*. **(G)** An *in vivo* xenograft model showed reduced tumor size when AKR1B1 KO A549 cells were used. **(H)** and **(I)** An *in vivo* xenograft model showed growth suppression when AKR1B1 silenced CY790 cells were used. **(J)** There was increased tumor mass when U87 cells were overexpressed in AKR1B1. *: *t*-test p<0.05; **: *t*-test p<0.01.

**Figure S5.**
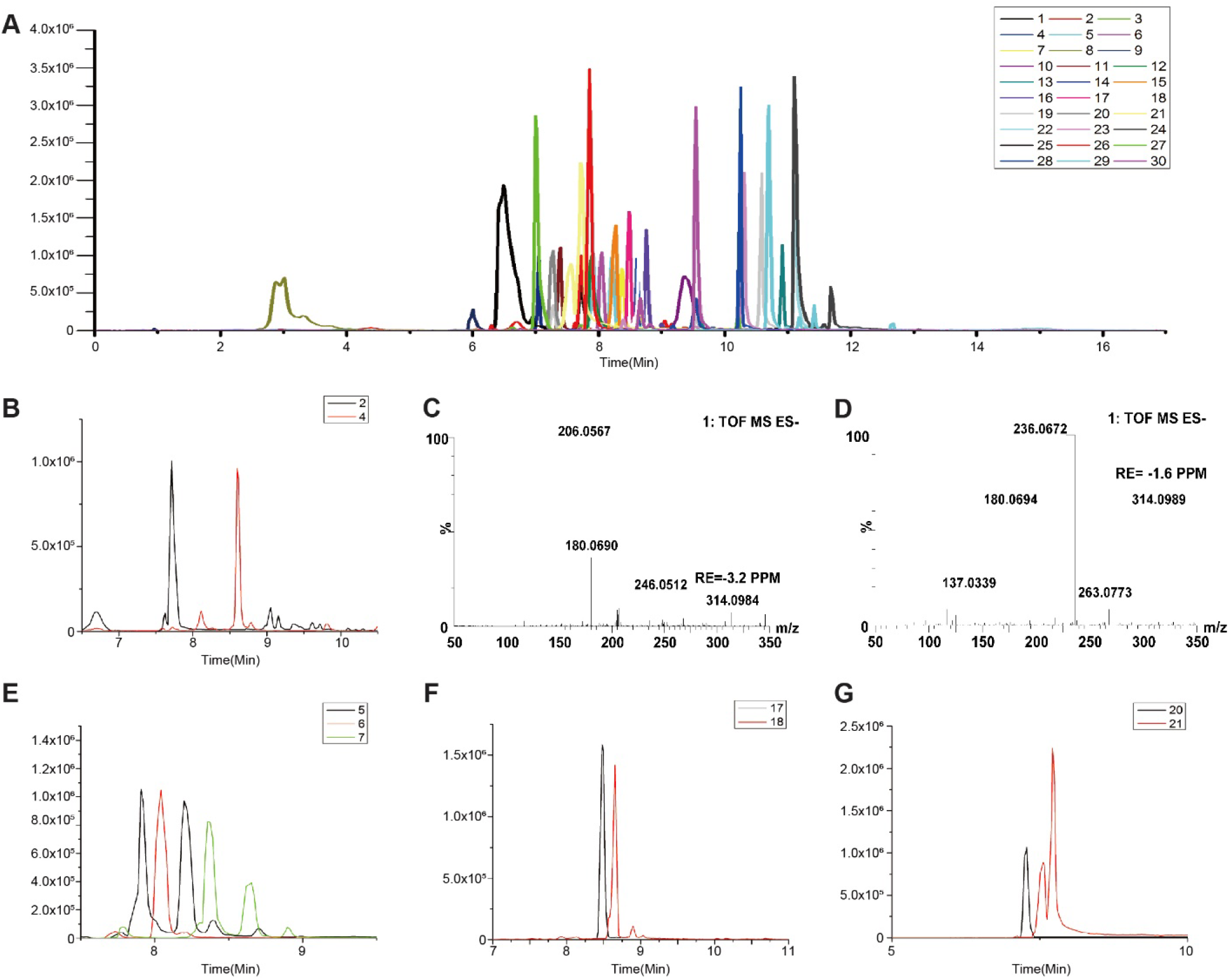
**(A)** UPLC/QTOF-MS of 3-NPH derivatives of metabolites. The labeled sugars are (1,2) D-Glucose; (3,4) D-Fructose; (5) Glucose 6-phosphate; (6) Fructose 1-phosphate; (7) Fructose 6-phosphate; (8) Glucose 1-phosphate; (9) Fructose 1,6-bisphosphate; (10) D-Glyceraldehyde 3-phosphate/Dihydroxyacetone phosphate; (11) 3-phosphoglyceric acid, and (12) Lactate; (13) Pyruvate; (14) 6-Phosphogluconic acid; (15) D-Sedoheptulose 7-phosphate; (16) D-Erythrose 4-phosphate; (17) D-Ribose 5-phosphate; (18) D-Ribulose 5-phosphate; (19) Succinic acid; (20) Isocitric acid; (21) Citric acid; (22) Oxoglutaric acid; (23) L-Malic acid; (24) Oxalacetic acid; (25) L-Serine; (26) Glutamic acid; (27) L-Glutamine; (28) Aspartic Acid; (29) Hydroxypyruvate; (30) Fumaric acid. **(B)** Representative IP-RP-UPLC-QTOF-MS chromatograms for separation (2)glucose and (4)fructose. **(C)** Mass spectrum of glucose obtained on QTOF mass spectrometry. **(D)** Mass spectrum of fructose obtained on QTOF mass spectrometry. Figure **(B)** and **(C)** show the postulated structures of the m/z 206.06 and m/z 236.07 fragment ions from glucose-3-NPHydrazone and fructose-3-NPHydrazone as a representative aldose and ketose, respectively. These two fragment ions were specific for aldoses and ketoses, respectively. The fragment ions m/z 206.06 and m/z 236.07 were therefore used for the qualitative monitoring of (1)glucose and (3)fructose. **(E)** Representative IP-RP-UPLC-QTOF-MS chromatograms for separation (5) Glucose 6-phosphate; (6) Fructose 1-phosphate; (7) Fructose 6-phosphate. **(F)** Representative IP-RP-UPLC-QTOF-MS chromatograms for separation (17) D-Ribose 5-phosphate; (18) D-Ribulose 5-phosphate. **(G)** Representative IP-RP-UPLC-QTOF-MS chromatograms for separation (20) Isocitric acid; (21) Citric acid.

**Table S1.**
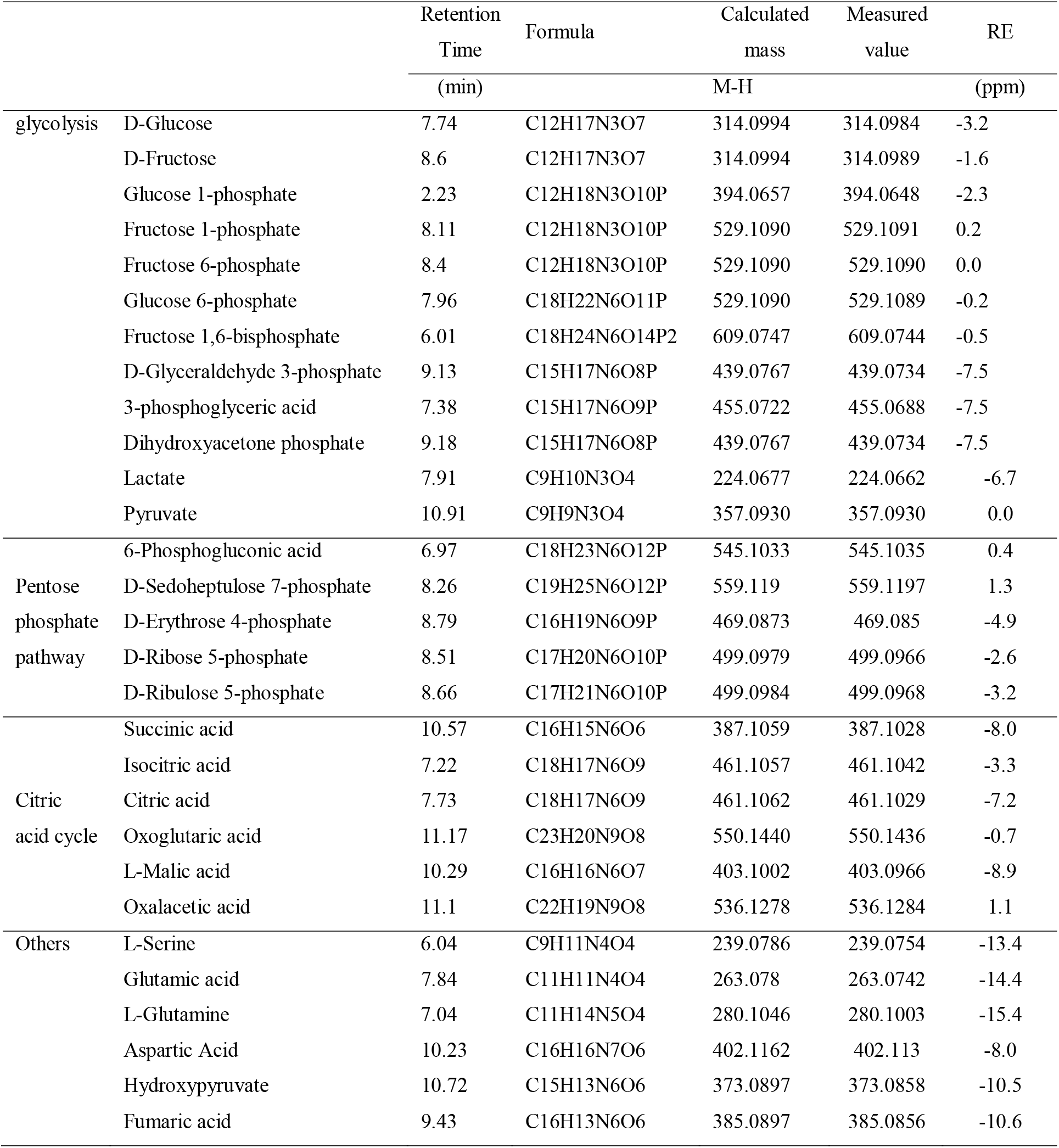
Identification of metabolites in cells using UPLC-QTOF-MS.

## References

Abdel-Haleem, A.M., Lewis, N.E., Jamshidi, N., Mineta, K., Gao, X., and Gojobori, T. (2017). The Emerging Facets of Non-Cancerous Warburg Effect. Front Endocrinol (Lausanne) 8, 279.

Amelio, I., Cutruzzola, F., Antonov, A., Agostini, M., and Melino, G. (2014). Serine and glycine metabolism in cancer. Trends Biochem Sci 39, 191–198.

Bononi, A., Giorgi, C., Patergnani, S., Larson, D., Verbruggen, K., Tanji, M., Pellegrini, L., Signorato, V., Olivetto, F., Pastorino, S., et al. (2017a). BAP1 regulates IP3R3-mediated Ca(2+) flux to mitochondria suppressing cell transformation. Nature 546, 549–553.

Bononi, A., Yang, H., Giorgi, C., Patergnani, S., Pellegrini, L., Su, M., Xie, G., Signorato, V., Pastorino, S., Morris, P., et al. (2017b). Germline BAP1 mutations induce a Warburg effect. Cell Death Differ 24, 1694–1704.

Chen, W.L., Jin, X., Wang, M., Liu, D., Luo, Q., Tian, H., Cai, L., Meng, L., Bi, R., Wang, L., et al. (2020). GLUT5-mediated fructose utilization drives lung cancer growth by stimulating fatty acid synthesis and AMPK/mTORC1 signaling. JCI Insight 5.

Chen, W.L., Wang, Y.Y., Zhao, A., Xia, L., Xie, G., Su, M., Zhao, L., Liu, J., Qu, C., Wei, R., et al. (2016). Enhanced Fructose Utilization Mediated by SLC2A5 Is a Unique Metabolic Feature of Acute Myeloid Leukemia with Therapeutic Potential. Cancer Cell 30, 779–791.

Debosch, B.J., Chen, Z., Saben, J.L., Finck, B.N., and Moley, K.H. (2014). Glucose transporter 8 (GLUT8) mediates fructose-induced de novo lipogenesis and macrosteatosis. J Biol Chem 289, 10989–10998.

Dong, Y., Yan, K., Ma, Y., Yang, Z., Zhao, J., and Ding, J. (2016). A modified LC-MS/MS method to simultaneously quantify glycerol and mannitol concentrations in human urine for doping control purposes. J Chromatogr B Analyt Technol Biomed Life Sci 1022, 153–158.

Duran, M., Beemer, F.A., Bruinvis, L., Ketting, D., and Wadman, S.K. (1987). D-glyceric acidemia: an inborn error associated with fructose metabolism. Pediatr Res 21, 502–506.

Goncalves, M.D., Lu, C., Tutnauer, J., Hartman, T.E., Hwang, S.K., Murphy, C.J., Pauli, C., Morris, R., Taylor, S., Bosch, K., et al. (2019). High-fructose corn syrup enhances intestinal tumor growth in mice. Science 363, 1345–1349.

Goto, Y., Hotta, N., Shigeta, Y., Sakamoto, N., and Kikkawa, R. (1995). Effects of an aldose reductase inhibitor, epalrestat, on diabetic neuropathy. Clinical benefit and indication for the drug assessed from the results of a placebo-controlled double-blind study. Biomed Pharmacother 49, 269–277.

Han, J., Lin, K., Sequria, C., Yang, J., and Borchers, C.H. (2016). Quantitation of low molecular weight sugars by chemical derivatization-liquid chromatography/multiple reaction monitoring/mass spectrometry. Electrophoresis 37, 1851–1860.

Jang, C., Hui, S., Lu, W., Cowan, A.J., Morscher, R.J., Lee, G., Liu, W., Tesz, G.J., Birnbaum, M.J., and Rabinowitz, J.D. (2018). The Small Intestine Converts Dietary Fructose into Glucose and Organic Acids. Cell Metab 27, 351–361 e353.

Jeremy M Berg, J.L.T., and Lubert Stryer. (2002). The Glycolytic Pathway Is Tightly Controlled. In Biochemistry (New York: W H Freeman).

Krause, N., and Wegner, A. (2020). Fructose Metabolism in Cancer. Cells 9.

Krzeslak, A., Wojcik-Krowiranda, K., Forma, E., Jozwiak, P., Romanowicz, H., Bienkiewicz, A., and Brys, M. (2012). Expression of GLUT1 and GLUT3 glucose transporters in endometrial and breast cancers. Pathol Oncol Res 18, 721–728.

Manolescu, A., Salas-Burgos, A.M., Fischbarg, J., and Cheeseman, C.I. (2005). Identification of a hydrophobic residue as a key determinant of fructose transport by the facilitative hexose transporter SLC2A7 (GLUT7). J Biol Chem 280, 42978–42983.

Ohkuma, T., Peters, S.A.E., and Woodward, M. (2018). Sex differences in the association between diabetes and cancer: a systematic review and meta-analysis of 121 cohorts including 20 million individuals and one million events. Diabetologia 61, 2140–2154.

Paglia, G., Hrafnsdottir, S., Magnusdottir, M., Fleming, R.M., Thorlacius, S., Palsson, B.O., and Thiele, I. (2012). Monitoring metabolites consumption and secretion in cultured cells using ultra-performance liquid chromatography quadrupole-time of flight mass spectrometry (UPLC-Q-ToF-MS). Anal Bioanal Chem 402, 1183–1198.

Quattrini, L., and La Motta, C. (2019). Aldose reductase inhibitors: 2013-present. Expert Opin Ther Pat 29, 199–213.

Scheepers, A., Schmidt, S., Manolescu, A., Cheeseman, C.I., Bell, A., Zahn, C., Joost, H.G., and Schurmann, A. (2005). Characterization of the human SLC2A11 (GLUT11) gene: alternative promoter usage, function, expression, and subcellular distribution of three isoforms, and lack of mouse orthologue. Mol Membr Biol 22, 339–351.

Sullivan, L.B., Gui, D.Y., Hosios, A.M., Bush, L.N., Freinkman, E., and Vander Heiden, M.G. (2015). Supporting Aspartate Biosynthesis Is an Essential Function of Respiration in Proliferating Cells. Cell 162, 552–563.

Sun, S.Z., and Empie, M.W. (2012). Fructose metabolism in humans - what isotopic tracer studies tell us. Nutr Metab (Lond) 9, 89.

Vander Heiden, M.G., Cantley, L.C., and Thompson, C.B. (2009). Understanding the Warburg effect: the metabolic requirements of cell proliferation. Science 324, 1029–1033.

Xu, S., Catapang, A., Doh, H.M., Bayley, N.A., Lee, J.T., Braas, D., Graeber, T.G., and Herschman, H.R. (2018). Hexokinase 2 is targetable for HK1 negative, HK2 positive tumors from a wide variety of tissues of origin. J Nucl Med.

Yabe-Nishimura, C. (1998). Aldose reductase in glucose toxicity: a potential target for the prevention of diabetic complications. Pharmacol Rev 50, 21–33.

